# Characterization of fluorescence lifetime of organic fluorophores for molecular imaging in the SWIR window

**DOI:** 10.1101/2022.12.16.520424

**Authors:** Luis Chavez, Shan Gao, Xavier Intes

## Abstract

**Significance:** Fluorescence lifetime imaging in the short-wave infrared (SWIR) is expected to enable high resolution multiplexed molecular imaging in highly scattering tissue.

**Aim:** To characterize the brightness and fluorescence lifetime of commercially available organic SWIR fluorophores and benchmark them against the tail emission of conventional NIR-excited probes.

**Approach:** Characterization was performed through our established Time-domain Mesoscopic Fluorescence Molecular Tomography (TD-MFMT) system integrated around a TCSPC-SPAD array. Brightness and fluorescence lifetime was measured for NIR and SWIR probes above 1000 nm. Simultaneous probe imaging was then performed to assess their potential for multiplexed studies.

**Results:** NIR probes outperformed SWIR probes in brightness while the mean fluorescence lifetimes of the SWIR probes were extremely short. The phantom study demonstrated the feasibility of lifetime multiplexing in the SWIR window with both NIR and SWIR probes.

**Conclusions:** Long tail emission of NIR probes outperformed the SWIR probes in brightness beyond 1000 nm. Fluorescence lifetime was readily detectable in the SWIR window, where the SWIR probes showed shorter lifetimes compared to the NIR probes. We demonstrate the feasibility of lifetime multiplexing in the SWIR window, which paves the way for in vivo multiplexed studies of intact tissues at improved resolution.

## 1 Introduction

Fluorescence optical imaging is an invaluable research tool which allows for non-invasive monitoring of numerous biological processes in live biological systems. Thanks to its high sensitivity, continuously growing library of available probes, and ability to image multiple probes simultaneously, fluorescence optical imaging techniques in the near-infrared spectral window (NIR) [700-1000 nm] have found numerous pre- and clinical applications.^1^ However, the resolution of these approaches is hampered by the scattering nature of tissues. As the scattering properties of tissue follow a power law in which the wavelength is the exponent, imaging further in the red has long been expected to improve imaging resolution. Only recently, thanks to the increasing availability of indium gallium arsenide (InGaAs) sensors with extremely high quantum efficiency (> 80 %) in the shortwave infrared (SWIR) spectral range [1000 – 2000 nm], far-red imaging in vivo has been demonstrated to be feasible at depths comparable to NIR methods. In conjunction with dedicated probes, emission SWIR imaging has been demonstrated to result in enhanced resolution,^2^ especially when using nanoparticles,^3^ while benefiting from lower tissue autofluorescence. However, such probes are limited to vascular imaging due to their sizes and SWIR dedicated organic fluorophores^4^ are required for enabling the vast majority of biological processes monitored by NIR probes. Alternatively, it has been proposed to acquire the long tail spectral emission of conventional near-infrared (NIR)-excited probes in the SWIR window.^5^ Still, such approaches are limited due to the need to use long pass filters that do not allow for spectral discrimination between molecules when multiplexed studies are involved. This drawback may be overcome using another distinctive fluorophore intrinsic property, its lifetime.

Fluorescence lifetime imaging (FLI) in the NIR window offers the ability to monitor intracellular parameters in preclinical applications,^6^ including cellular metabolism^7^ and pH,^8^ as well as to quantify Förster resonant energy transfer (FRET) for monitoring probe target engagement.^9, 10^ Integrating FLI into well-established intensity-based imaging systems can complement their performances; recently, our group characterized and validated scanning-based Time-domain Mesoscopic Fluorescence Molecular Tomography (TD-MFMT), capable of acquiring fluorescence intensity for 3D probe reconstruction as well as resolving fluorescence lifetime information in the visible-NIR window.^11^ Nevertheless, challenges in implementing FLI in SWIR arise from limitations in both SWIR-specific probes and detectors. Current studies mostly use inorganic probes that can be used for luminescence lifetime multiplexing^12^ but which raise biocompatibility issues.^13^ Besides, intensity in the long tail emission of NIR probes represent only a small fraction of the maximum intensity, requiring detectors with high quantum efficiency (QE) and low dark current noise for effective signal-to-noise (SNR) detection.^2^ Moreover, the fluorescence lifetime of organic NIR fluorophores decrease as the wavelength approaches the SWIR window,^14^ demanding systems with high temporal sensitivity for lifetime quantification. Furthermore, the commonly used time-resolved detectors for lifetime quantification encounter difficulties to extend their utilities in the SWIR range. Beyond the drawbacks of mechanically fragile feature and large form factor, there are few photomultiplier tubes (PMTs) commercially available in the NIR, and even less in the SWIR.^15^ Another preferred imager in the NIR, the gated CCD, has its QE cut-off above 900 nm and hence hinders its implementation for longer wavelength detection. Alternatively, due to the relatively decent QE in the SWIR range and the impressive temporal sensitivity, silicon-based single-photon avalanche diode (SPAD) array^16^ coupled with a time-correlated single-photon counting (TCSPC) module has the potential to quantify the short lifetime of the SWIR-specific fluorophores. Herein, we report on the brightness and lifetime characteristics of ubiquitous NIR and SWIR fluorophores using such detector.

Previous research has characterized the brightness of commercially available NIR and SWIR-specific probes in the NIR-SWIR windows^2, 5^ and the lifetime of organic NIR fluorophores has been well documented.^13^ However, there is still a lack of studies characterizing the lifetime of commercially available SWIR fluorophores. Herein, we report for the first time the lifetime characterization of SWIR-specific fluorophores IR-E1050 and IR-T1050. Additionally, we compare both brightness and lifetime of the SWIR fluorophores against the tail emission of conventional NIR fluorophores – IRDye800CW and Alexa Fluor 750 in the SWIR window by leveraging TD-MFMT system in well-plate settings. By performing phantom experiments, we demonstrate the lifetime multiplexing in the SWIR regime by distinguishing short-lifetime fluorophores and accurately quantifying the relative concentration fraction of two mixed dyes.

## 2 Methods and Results

To characterize the performance of NIR/SWIR fluorophores in regards to brightness and lifetime properties, we employed our established TD-MFMT system to leverage its outstanding temporal sensitivity (up to ~ 90 ps of full width at half maximum (FWHM) of the instrument response function (IRF) and 1.6 ps time step resolution). For the NIR dyes, we selected IRDye 800CW (LI-COR Bioscience) and Alexa Fluor 750 (AF750, ThermoFisher) due to their ubiquitous use in the field. For the SWIR-specific fluorophores, we employed commercially available probes, IR-E1050 and IR-T1050 (Nirmidas Biotech).^2^ Compared to the diagram reported previously,^11^ TD-MFMT system was adapted by replacing the sCMOS with the InGaAs camera (Owl 640 II, Raptor Photonics) and switching the optics to the SWIR-optimized components (shown in Fig. 1) for optimal detection efficiency. A wide-field (WF) illumination was additionally set up for acquiring WF intensities by the InGasAs camera for brightness comparison. Sequentially, with the scanned point illumination, TD data sets were obtained by the SPAD array coupled with the TCSPC module for lifetime quantification. A tunable Ti:Sapphire pulse laser (MaiTai, Spectral Physics) is used by the WF acquisition as well as the scanning-based TD measurements producing pulses across 690 nm to 1040 nm.

**Fig 1.**
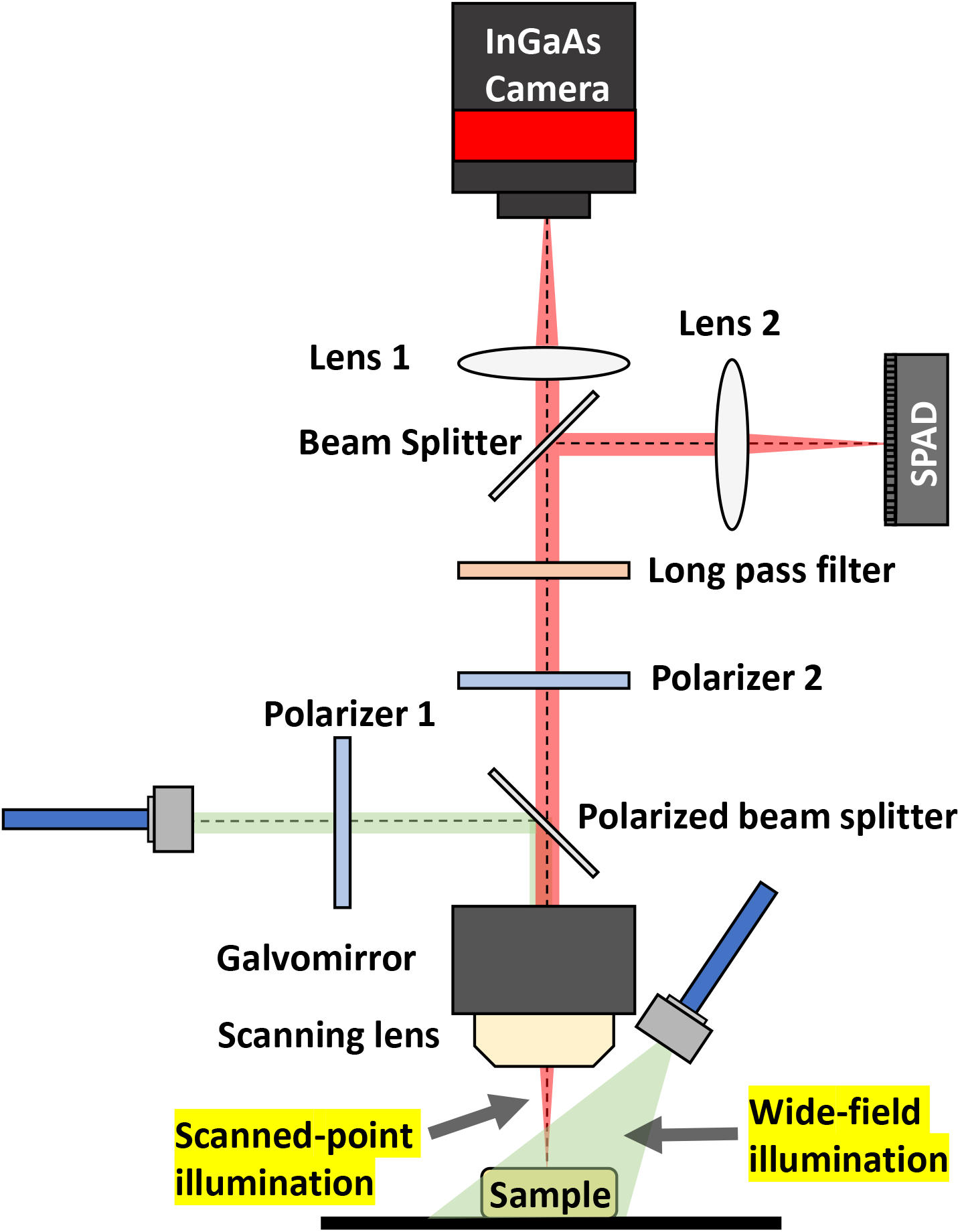
Adapted TD-MFMT system schematic for SWIR lifetime characterization

The four fluorophores were diluted in phosphate-buffered saline (PBS) solution to a concentration of 5 μM for well-plate and phantom imaging experiments. With the exception of IRDye800CW, which was excited using 775 nm wavelength, the rest three fluorophores were excited at 745 nm, and all four fluorophores were imaged with the same long-pass emission filter (FELH1000, Thorlabs). The laser power was set as 6.5 mW/cm^2^ and 62 mW/cm^2^ for WF and TD acquisitions respectively (well below the Maximum Permissible Exposure). Following by a single shot of WF acquisition, temporal point spread functions (TPSFs) were captured at each scanning position by the SPAD and fed into a mono-exponential fitting model for the pixel-wise lifetime quantification using the open source software AlliGator.^17, 18^ One can follow the workflow of the TD data collection and lifetime quantification laid out in our previous work.^11^ WF and SPAD-based intensities measured by the InGaAs camera and the SPAD, the lifetime quantification results, as well as the normalized IRF and representative decays of four fluorophores are shown in Fig. 2 A and C. Figure 2 B illustrates the pixel-wise SPAD-based intensity distribution of the four fluorophores. Our results show that when excited at their optimal absorption wavelength, the tail emission of the NIR-excited fluorophores emitted a brighter signal compared to the SWIR fluorophores, where signal from IRDye800CW and AF750 was approximately four and three times brighter than the brightest SWIR probe, respectively. Furthermore, the characterization of the mean lifetime of the probes indicates the SWIR probes exhibiting very short lifetimes, confirming the aforementioned relationship between emission wavelength and fluorescence lifetime. These findings are summarized in Table 1.

**Fig 2.**
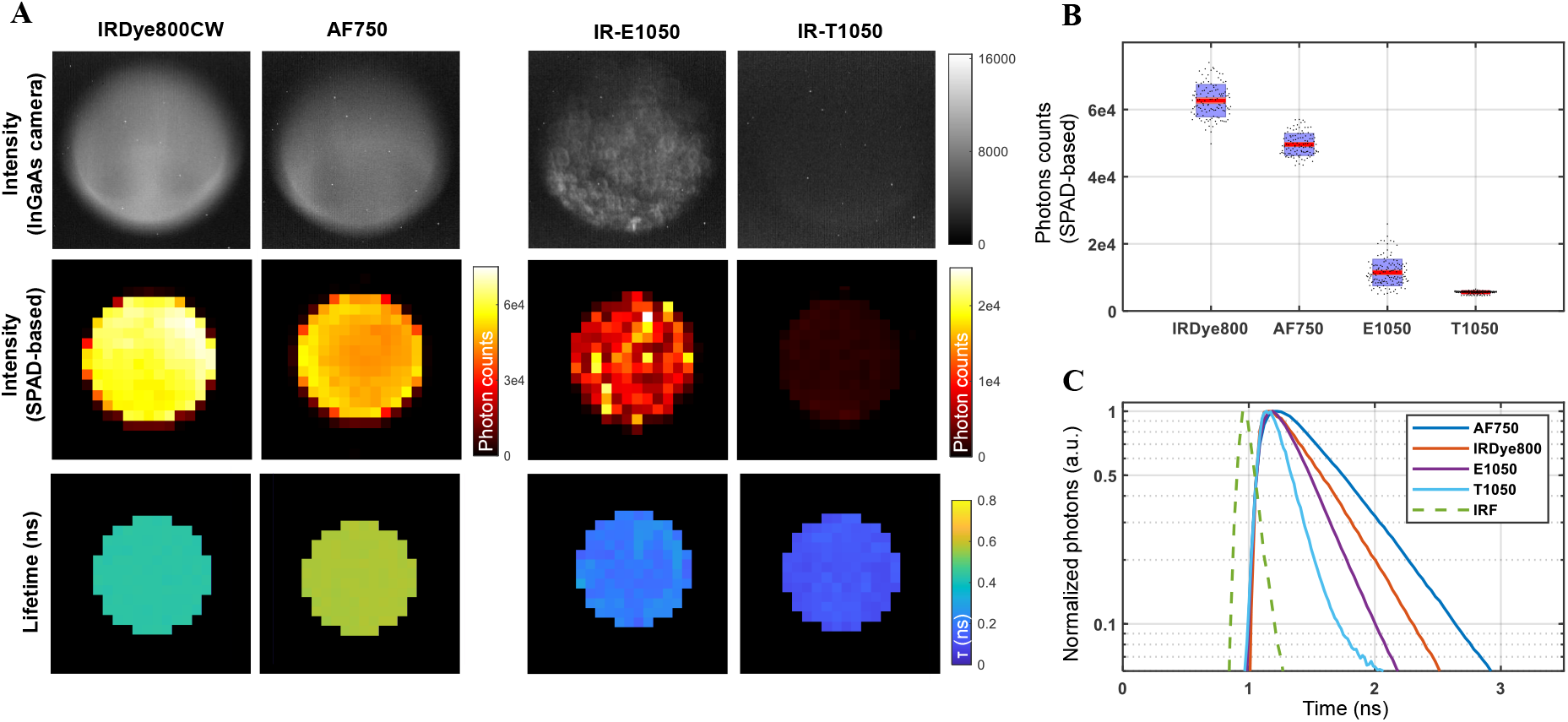
Intensity comparison and lifetime quantification results with well-plate settings. A: WF/SPAD-based intensity and lifetime maps for four fluorophores (scanning point = 20 × 20, scanning step = 250 μm, Scale bar = 1 mm). B: Pixel-wise photon count distribution measured with SPAD in box plot. C: Normalized representative fluorescence decays and IRF in logarithmic scale.

**Table 1.**
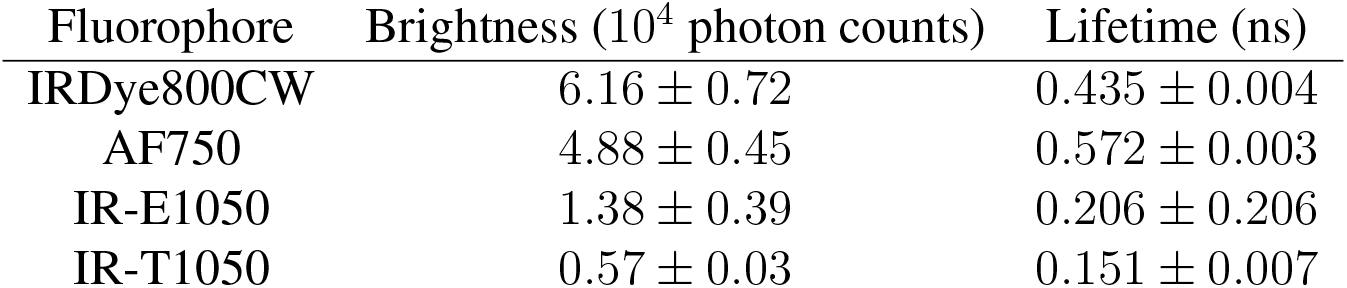
Summary of fluorophore characterization (mean ± standard deviation)

**Table 2.**
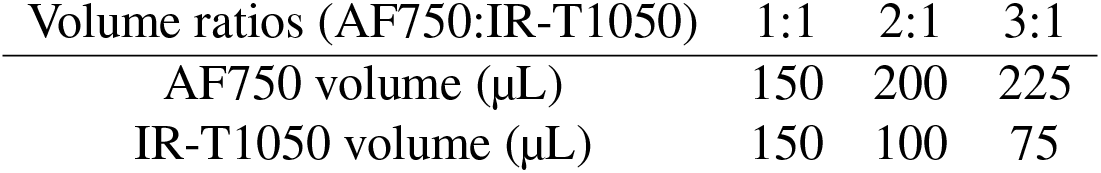
Volumes of fluorescence dyes for lifetime unmixing

| Volume ratios (AF750:IR-T1050) | 1:1 | 2:1 | 3:1 |
| --- | --- | --- | --- |
| AF750 volume ($\mu\text{L}$) | 150 | 200 | 225 |
| IR-T1050 volume ($\mu\text{L}$) | 150 | 100 | 75 |

To validate the potential for multiplexed imaging using lifetime contrast while acquiring in the SWIR spectral range, a phantom was 3D printed with RPI letter grooves (paradigm shown in Fig. 3 A), which contains AF750, IRDye800CW and IR-E1050 at 5 μM for letters “R”, “P” and “I”, respectively. The illumination wavelength was tuned at 750 nm for maximizing the absorption efficiency of all the fluorophores. WF and TD data sets were captured and processed following the same imaging protocol in the well-plate experiments, though the whole sample was imaged at once using the same system settings. The results shown in Fig. 3 establish the feasibility of simultaneous detection of multiple probes in the SWIR window through multiple modalities at clinically relevant concentrations and with the same excitation setting. Intensity detection shown in Fig. 3 B and C demonstrates that probe distribution can be readily monitored at high resolution, despite significant variations in brightness between different probes. Similarly, fluorescence lifetime quantification results shown in Fig. 3 D illustrate the ability to quantify lifetime values accurately while using our SPAD-array as well as significant lifetime contrast between probes. To demonstrate the potential of using such lifetime contrast for multiplexing studies, we mixed AF750 and IR-T1050 together at different concentration ratios. 3 μM AF750 and 100 μM IR-T1050 solutions were prepared to obtain comparable intensities while using the same excitation wavelength (750 nm laser) and emission settings (long-pass filter FELH1000) aforementioned. Three wells were filled with both dyes to get a total volume of 300 μL at different volume ratios ranging from 1:1 to 3:1 (AF750:IR-T1050). TD imaging was conducted following the same protocol as previously described and the data sets were fitted using a bi-exponential decay model in AlliGator with initial lifetimes (*τ*_*AF*750_ = 0.57 ns and *τ*_*T*1050_ = 0.15 ns) based on the prior mono-exponential fitting results. The SPAD-based intensity maps, the amplitude fraction maps of AF750 as well as the representative TPSFs with different volume fractions are illustrated in Fig. 4 A and C. The amplitude fraction was extracted from the fitting and can be correlated with the fluorophore fraction in the mixture, where the amplitude fraction of AF750 tends to increase as the volume of AF750 becomes dominant in the mixture (shown in Fig. 4 B), thus demonstrating the potential of lifetime as a contrast mechanism to unmix fluorophores in the SWIR window while using long pass filter for optimal brightness.

**Fig 3.**
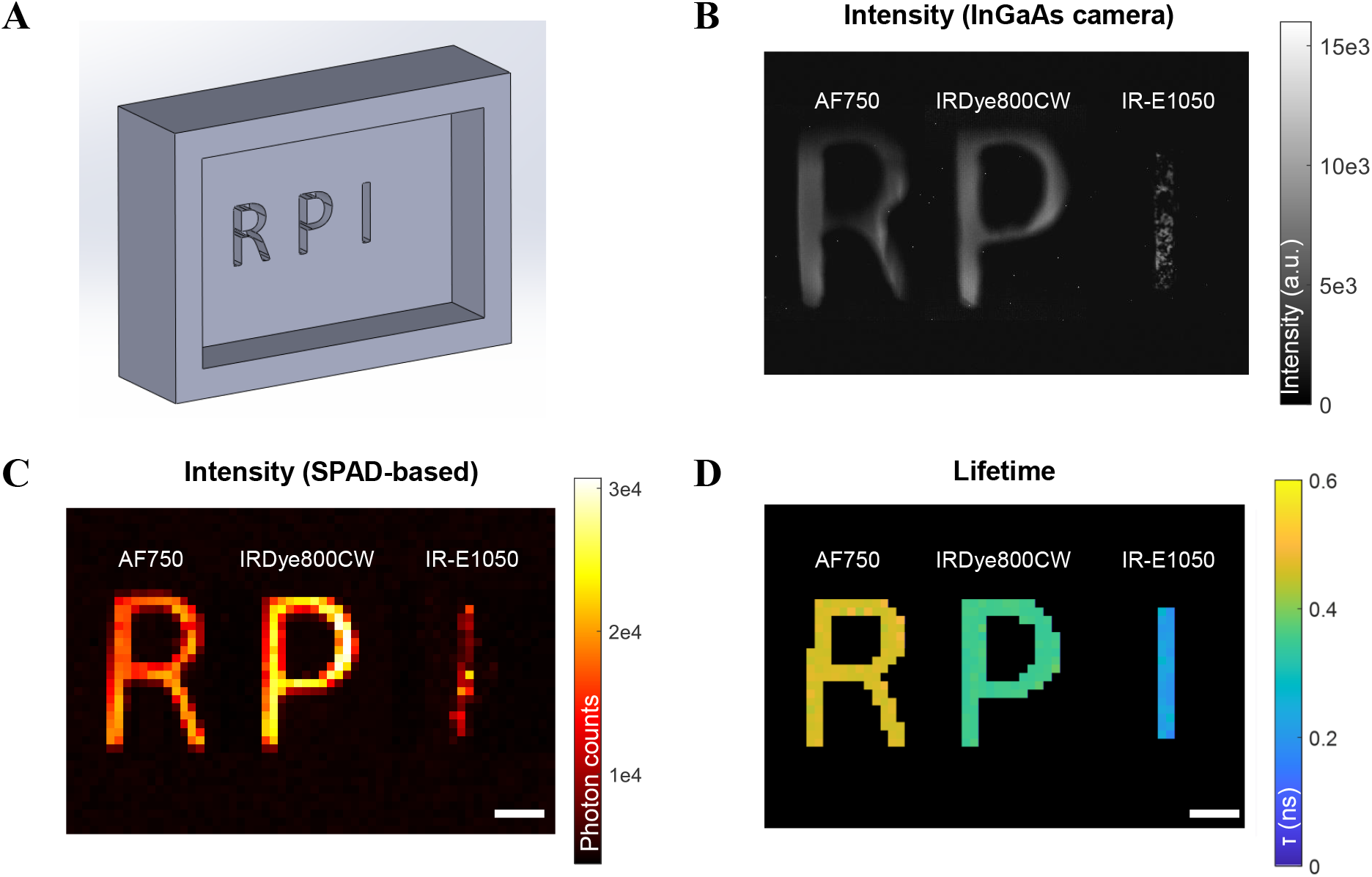
Lifetime quantification by liquid phantom experiment. A: 3D-printed RPI phantom paradigm. B, C: WF/SPAD-based intensity. D: Lifetime map. (scanning point = 40 × 60, scanning step = 250 μm). Scale bar = 1 mm

**Fig 4.**
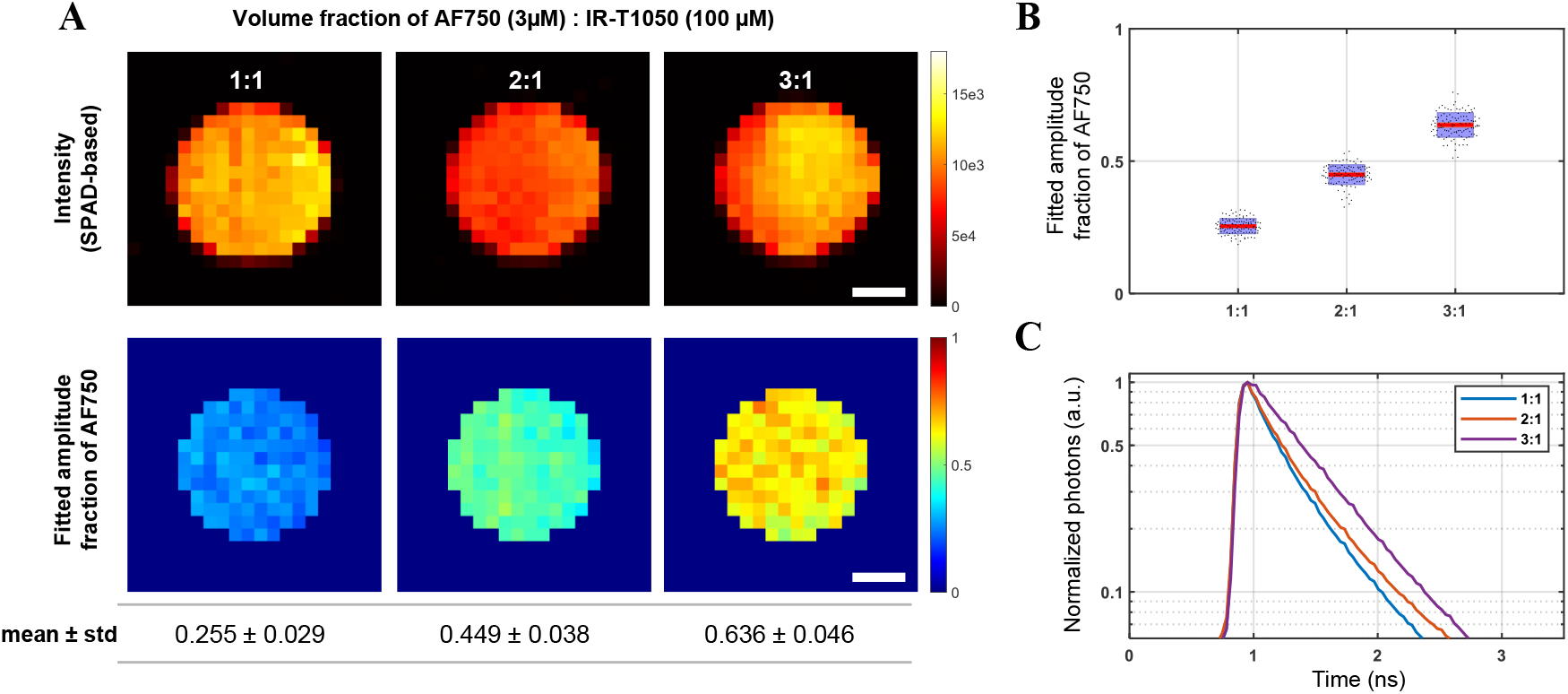
Lifetime multiplexing by two fluorescence dye. A: SPAD-based intensity map and fitted amplitude fraction of AF750. B: Pixel-wise distribution of fitted amplitude fraction of AF750 in box plot. C: Representative fitted decays. (scanning point = 20 × 20, scanning step = 250 μm). Scale bar = 1 mm

## 3 Discussion

Concurrent monitoring of probe intensity and lifetime in the SWIR window promises to enhance current in vivo optical molecular imaging applications at the mesoscopic and macroscopic scales by combining gain in imaging resolution to better resolve the biodistribution of probes with monitoring multiple processes or additional molecular information via lifetime sensing. However, limitations in detection hardware and lack of suitable SWIR probes remain a challenge in the translation from NIR to SWIR imaging. Herein, we report a quantitative comparison between brightness and lifetime measurements of two conventional NIR probes and two commercially available SWIR probes in the SWIR imaging window. Fluorescent signal for all probes was detectable at clinically relevant concentrations, where the NIR probes yielded a higher intensity compared to the SWIR probes. Although a direct comparison between the brightness of IRDye800CW and IR-E1050 has been previously reported showing IRDye800CW being brighter in the SWIR window,^2^ we show that NIR probe AF750 outperforms the SWIR fluorophores in brightness, noting that its peak emission is blue shifted compared to that of IRDye800CW.

It is noteworthy that fluorescent temporal decays were readily acquired beyond 1000 nm with good SNR given the still relative low quantum efficiency of our SPAD array in the SWIR window (~5%). The high photon counts acquired on relatively low concentration samples related with the short IRF associated with the SPAD array (~120 ps) enabled to quantify lifetimes with accuracy. To our knowledge, this is the first report on the lifetimes of these two SWIR fluorophores. Characterizing the fluorescent lifetime of the SWIR probes further shows the current superiority in applicability of the NIR probes. Shorter fluorescence lifetime requires detectors with higher temporal sensitivity for effective detection as well as increased experimental control to ensure temporal stability (reduce drift and jitter). Moreover, for in vivo applications, shorter lifetimes estimation can be affected by the topography of the boundary condition as well as the depth of the fluorescence inclusion. Hence, quantifying short lifetime robustly is more difficult at large and requires increased expertise.

Overall, these results complement previous characterization of conventional NIR fluorophores. It has been previously demonstrated that these fluorophores emit fluorescent signal well into the SWIR window similarly to the ones presented herein, where signal is detected even when excited at a non-optimal wavelength.^5^ We extended further this work by demonstrating effective detection of fluorescent lifetime with a SPAD array optimized for measurements in the VIS-NIR window. In turn, this lifetime information was used for quantifying fluorophore fraction of well controlled mixtures. These results pave the way for lifetime multiplexing in the SWIR window using the vast library of conventional NIR fluorophores and single excitation wavelength, which should provide invaluable information in preclinical studies until suitable SWIR fluorophores become available.

In summary, the experimental results show that the conventional NIR-excited fluorophores out-performed the commercially available SWIR probes in both brightness and lifetime in the SWIR window. Furthermore, effective detection of both lifetime and intensity of tail emission in the SWIR window shows promise for the translation of multimodal fluorescence imaging into the SWIR region with the current library of NIR fluorophores. Hence, this study demonstrates the feasibility of lifetime multiplexing in the SWIR range using the tail emission of conventional fluorophores over current commercially available organic SWIR-specific probes.

## Acknowledgments

This work was supported by the following grants from the National Institute of Health (R01CA207725, R01CA237267, and R01CA250636).

## Code, Data, and Materials Availability

All analyses are performed using freely available software.^17^ Data underlying the results presented in this paper are not publicly available at this time but may be obtained from the authors upon reasonable request.

**Xavier Intes** received his PhD from the Université de Bretagne Occidentale and a postdoctoral training at the University of Pennsylvania. He is a professor in the Department of Biomedical Engineering, Rensselaer Polytechnic Institute, and an AIMBE/SPIE/Optica fellow. He acted as the chief scientist of Advanced Research Technologies Inc. His research interests are on the application of diffuse functional and molecular optical techniques for biomedical imaging in preclinical and clinical settings.

Biographies and photographs of the other authors are not available.

